# The influence of gene *Rv3671c* in *Mycobacterium bovis* to its replication and acid resistance

**DOI:** 10.1101/2020.09.15.298273

**Authors:** Weidong Lin, Ting Xin

## Abstract

To study the effect of Marp protein in Mycobacterium *bovis* to the acid resistance and growth performance, this research constructed a knockout mutant (*ΔMarp*) with mycobacteriophage, complemented strain (ΔMarpComp), and overexpressing strain (PmvRv3671) of the gene *Rv3671c* with pmv261 plasmid. Culturing them in standard 7H9 medium to the early logarithmic phase and transferring them into 7H9 medium at pH 6.6 and pH 5.0 and maintenance solution at pH 6.6 and pH 4.5. Likewise, macrophages Raw264.7 were infected with multiple infections at 10. The results showed that while they grew in 7H9 medium at pH 5.0 or pH 6.6 and maintenance buffer at pH 4.5, the lived number of over-expressing strain PmvRv3671 is more than wild-type strain M.bovis, ΔMarpComp and *ΔMarp* on the 14^th^ day. After removing the effects of citrate solution, it can be found that the acid resistance abilities of them all are significantly lower on the 14^th^ day than that on the 5^th^ day. Using them infected Raw264.7 macrophages with IFNγ stimulation, the growth rate of the PmvRv3671 is better than An5, *ΔMarp* and ΔMarpComp. In conclusion, *Rv3671c* over-expressing strain had shown a better growth ability than wild-type An5, *ΔMarp* and ΔMarpComp under acidic environment. When exposed to pH 4.5 citrate maintenance solution for a long time, acid resistance abilities of them all have become weaker.

## 1 Introduction

Tuberculosis is an important infectious disease with large number of deaths caused by a single pathogen in the world and is also considered the leading cause of death after AIDS (WHO, 2019).

The main pathogenic bacteria of tuberculosis are Mycobacterium tuberculosis (Mtb) and Mycobacterium bovis (*M. bovis*). Mechnikov indicated that macrophages would kill most of the ingested microorganisms by acidifying its interior, but Mtb could use its waxy cell wall to resist this acidification (Mechnikov, 1988). According to previous reports, when Bacille Calmette-Guérin (BCG), an attenuated variant of *M. bovis*, was exposed to pH 5.5 in vitro, it ceased replicating and Mtb continued to divide slowly unless the pH was lowered to 4.5. With the addition of 0.5 mM nitrite to 7H9 medium at pH 5.5, Mtb would also stop replicating. However, it is in a balanced state, where the Mtb could survive for several days, and the bacterial can return to a normal growth rate after replacing the medium with standard 7H9 (Bryk et al., 2008; Gold et al., 2012; Vandal et al., 2008). Based on that, Vandal used the acidic environment to screen 10,100 MTB transposon mutants and found a mutant that cannot maintain its intracellular pH homeostasis under a pH 4.5 environment (Vandal et al., 2008). Because the gene *Rv3671c* product played an essential role in the acid resistance of Mtb, it is named as Mycobacterial acid resistance protease (Marp). Marp was identified as a transmembrane serine peptidase with a protease domain located in the periplasm, as confirmed by a prudent analysis of its mutations, homology modelling, crystallography, and substrate profiling (Biswas et al., 2010; Small et al., 2013; Vandal et al., 2008).

We had confirmed that Marp was also presented in *M. bovis* (data not shown), but it remains as speculation if the role of this gene in *M. bovis* is related to acid resistance or if there are other biological functions involved in, such as the growth. Furthermore, because *M. bovis* had more extensive hosts beyond human susceptibility, it could also infect domestic animals and wild animals under natural circumstances (Pesciaroli et al., 2014). Thus further studies on *M. bovis* are extremely meaningful to public safety (Gao et al., 2019; Xin et al., 2018).

In order to study the functions and biological characteristics of the gene *Rv3671c* in *M. bovis*, this research aimed to construct a knockout mutant (*ΔMarp*), complemented strain (ΔMarpComp), and overexpressing strain (PmvRv3671) of the gene *Rv3671c*. After that, they were cultured in standard or acidic mediums and used to infect macrophages to closely examine and compare their difference. Subsequently, it was possible to state the functions and influences of gene *Rv3671c* in M. *bovis* to its growth and acid resistance.

## 2 Materials and methods

### 2.1 Analyzing Rv3671c sequence and designing the primers

According to the genome sequence of *M. bovis* (AWPL01000069.1) registered in GenBank, the search for the *Rv3671c* sequence showed that there is a high homology sequence between 158384 and 159586, consistent with the characteristics of *Rv3671c* (CP003248.2) in MTB H37Rv. As shown in Figure 1, the red band of the *Rv3671c* gene was knocked out of *M. bovis*. The blue bands of the upstream and downstream fragments of the gene are called the left arm and the right arm individually. The primers LFP / LRP and RFP / RRP are used to amplify the left and right arms. The primers LRP and RFP need to contain the partial sequence of the left and right ends of the gene to be eliminated. In this case, the verification primer (LYZ / RYZ) is used in the subsequent PCR verification stage, and the verification primer is designed upstream of the LFP and downstream of the RRP (indicated by the black band). The primer sequences are all listed in Table 1.

**Table 1.**
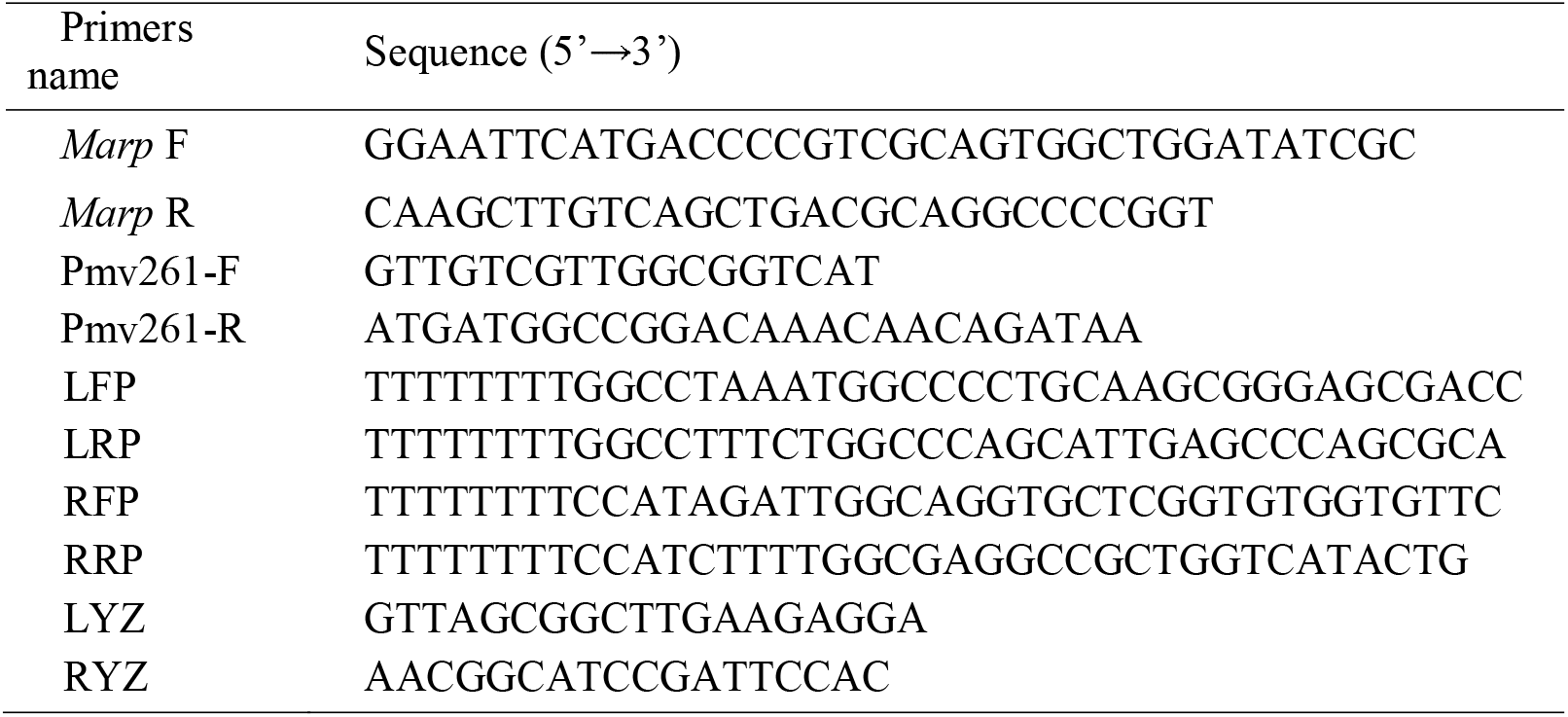
Primers used in this study.

**Figure 1.**
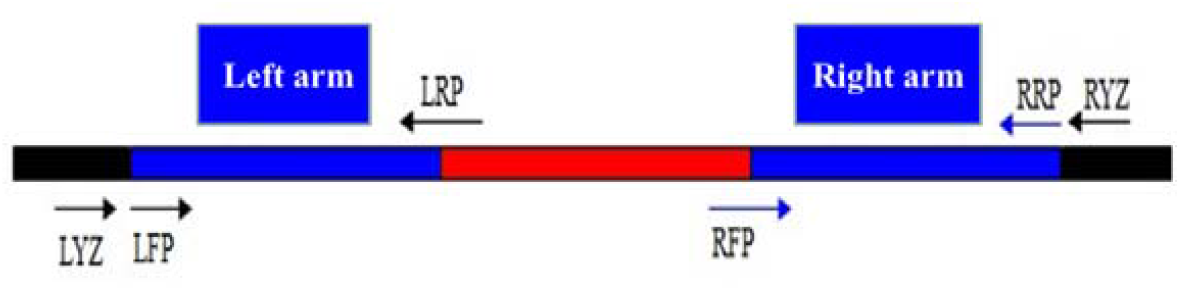
Schematic diagram design of primers. The blue bands of the upstream and downstream fragments of the gene are called the left arm and the right arm individually. LFP / LRP and RFP / RRP are used to amplify the left and right arms. The primers LRP and RFP need to contain the partial sequence of the left and right ends of the gene to be knocked out. The verification primer (LYZ / RYZ) is used in the subsequent PCR verification stage, and the verification primer is designed upstream of the LFP and downstream of the RRP (indicated by the black band).

### 2.2 Construction of M. bovis Rv3671c knockout mutants

For this chunk of the experiment, native *Rv3671c* was replaced in An5 *M. bovis* with a hygromycin-resistant cassette via allelic exchange following the protocol as reference (Bardarov et al., 2002). The DNA segments flanking upstream and downstream regions of the *Rv3671c* gene (named left arm and right arm) were amplified by PCR using suitable primers (as shown in Figure 2) and digested by *SrfI* and *PflMI* separately. The allelic exchange substrate was then cloned into the suicidal vector p0004S directionally, on either side of the hygromycin resistance-sacB gene cassette (3.6Kb). The recombinant vector p0004S was also linearized by *PacI* digestion, and packaged into the temperature-sensitive TM4 shuttle phasmid phAE159 with digesting by *PacI* digestion for generating the specialized transducing mycobacteriophage. The phage could be amplified in *M. smegmatis* mc^2^155, and high-titer phage particles were then prepared at a replication permissive temperature of 30°C. Further on, the gene was deleted from *M. bovis* An5 by specialized transduction. Allelic exchange occurred as a result of a double crossover between the homologous DNA arms flanking the disrupted gene. Transductants were then screened for gene disruption by PCR using a forward primer for left arm (LFP) and right arm reverse primer (RRP). Gene deletion could be confirmed by western blotting using monoantibody against Marp. The Rv3671c gene disruption strain was designated *ΔMarp*.

**Figure 2.**
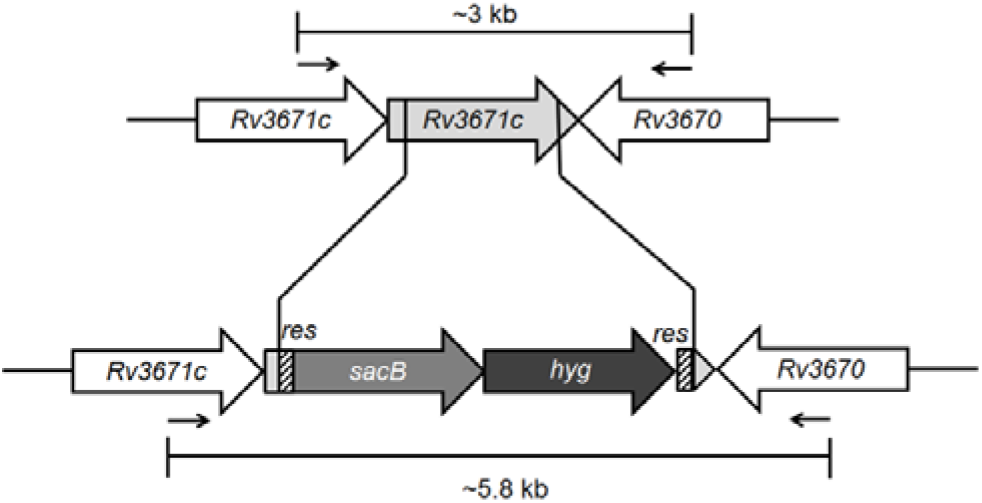
Schematic diagram of allelic replacement of the Rv3671c gene. The constructed phagemid contains the DNA segments flanking upstream and downstream regions of the Rv3671c gene (named left arm and right arm). So when the packaged phage infected with *M. bovis*, native Rv3671c was replaced in *M. bovis* An5 with a hygromycin-resistant cassette via allelic exchange.

### 2.3 Construction of complemented strain and overexpressing strain

For completing the *ΔMarp* and preparing an overexpressing strain, the *Rv3671c* gene was amplified from *M. bovis* genomic DNA by PCR. After that, it was cloned into the *E. coli*-*Mycobacterium* shuttle plasmid pMV261 with the primer-incorporated *HindIII* and *EcoR I* sites to yield the recombinant plasmid named pMV261-Marp. This recombinant plasmid was then electroporated into the mutant strain *ΔMarp* and wild-type strain An5 with 2.5 kV, 1,000 ohms and 25μF. Electroporation was performed as previously described (Snapper et al., 1990). The complemented colonies were selected from plates containing hygromycin and kanamycin, and overexpressing strains were selected from plates containing kanamycin. PCR and western blot were used to identify the genome and proteins of selected ΔMarpComp and PmvRv3671.

### 2.4 Media and Culture Conditions

*M. bovis* was cultured at 37 °C on Middlebrook 7H10 agar containing 10% oleic acid–albumin-dextrose-catalase (OADC) and 0.5% glycerol or in 7H9 standard medium (Middlebrook 7H9 broth supplemented with 0.2% glycerol, 10% OADC, and 0.05% Tween 80 or 0.02% Tyloxapol). The standard 7H9 was acidified to pH 4.5 or pH 5.0 with 2 N HCl (*M. bovis* couldn’t grow in 7H9 at pH4.5). According to the previous report, at least two components of the 7H9 medium can become toxic to MTB at pH 4.5: one associated with Tween and another associated with albumin, and both may release free fatty acids at low pH (Vandal et al., 2008). So the experiment needed to be supplemented with culturing these bacteria on 7H9 at pH5.0. Phosphate-citrate (pcit) buffer at pH 4.5 or pH 6.6 was made from 200mM sodium phosphate and 100 mM citric acid. Pcit buffer at pH 4.5 or pH 6.6 contained 0.02% tyloxapol (pcit-6.6 buffer or pcit-4.5 buffer). Strains with antibiotic resistance cassettes growled in the presence of 75 μg/mL hygromycin B (*ΔMarp*), 20 μg/mL kanamycin (PmvRv3671), or a combination of 75 μg/mL hygromycin B and 20 μg/mL kanamycin (COMP) (van Kessel and Hatfull, 2007).

### 2.5 Measurement of acid resistance and growth characters

Colony-forming units (CFU) were determined by plating serial dilutions of the suspensions on 7H10 agar plates. The early–log-phase (within 3days) cultures were washed with 7H9- pH6.6, 7H9- pH5.0, 7H9- pH4.5, pcit-ty-6.6 or pcit-ty-4.5 medium and centrifuged at 120g for 10 min. Also, the single-cell suspensions were adjusted to ∼5 × 10^6^ CFU/mL (∼OD_600_ = 0.015) in 7H9-pH6.6, 7H9-pH4.5, pcit-ty-6.6 or pcit-ty-4.5 medium and incubated at 37 °C. The pH and OD_600_ values were also recorded for further evaluation.

### 2.6 Intracellular viability of the M. bovis

The intracellular viability of *ΔMarp* was determined using macrophage infection RAW264.7 cells, cultured in DMEM medium supplemented with 10% fetal bovine serum (FBS) and 10 mM HEPES. The cells were then seeded onto 24 well plates (10^6^ cells/well) with or without 10 ng/mL murine IFN-γ (R&D Systems). Sixteen hours later, macrophages were used to infect in triplicates with wild type AN5, *ΔMarp*, PmvRv3671, and the ΔMarpComp at a multiplicity of infection of 0.1 and then washed with PBS 4 hours later to remove the non-phagocytosed bacilli. The infected macrophages were then incubated with fresh DMEM supplemented with 2% FBS at 37°C in the presence of 5% CO2 and it was necessary to replace the medium every 48 h. Intracellular bacilli could be recovered by lysing infected macrophages with 0.5% Triton X-100 and enumerated bacteria by plating serial dilutions of the lysate on 7H10 or 7H11 agar plates in 48, 96, and 144hours post-infection.

### 2.7 Statistical analysis

GraphPad Prism 5 software (GraphPad Software, Inc., USA) or SPSS software version 20.0 (IBM, Inc., USA) was used for the data analysis and all the data expressed as the mean ± standard error of the mean. Differences in the survival rates and growth ratios, between the different groups (An5, *ΔMarp*, ΔMarpComp, and PmvRv3671) were determined by two-way ANOVA and considered statistically significant at *p*<0.05 or highly significant at *p*<0.01.

## 3 Results

### 3.1 Amplifying the upstream and downstream homology arms of the Rv3671c gene

The amplification results of the upstream and downstream homology arms of the target gene by PCR are displayed in Figure 3a, and the sizes of PCR amplification products are consistent with the expected length. The upstream and downstream of homologous arms were digested by restriction enzymes *SrfI* and *PflMI*, respectively, and the digested products were recovered. After connecting the upstream and downstream to the vector p0004s, named p0004S-*ΔMarp*, it was transformed into *E*.*coli* DH5α. After culturing them on LB agar with hygromycin B, clones were picked up, and their plasmids were identified by PCR. The P0004S-*ΔMarp* plasmid was then packaging into phAE159 and transformed into HB101 competent and finally the plasmid was extracted from positive colonies by digestion identification (the results are shown in Figure 3b).

**Figure 3.**
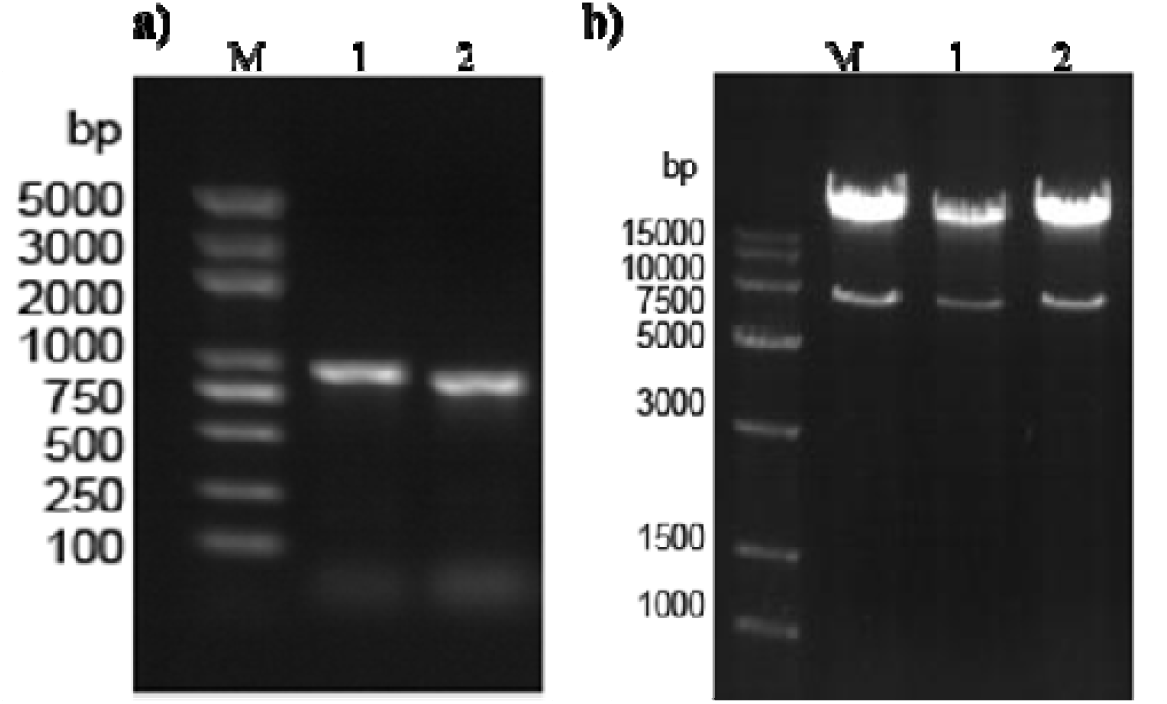
The process to build recombinant phAE159. a): PCR amplification results of the upstream and downstream homology arms of the Rv3671c gene. Extracting the DNA from *M. bovis* AN5, the template was used to amplify the upstream (lane 1) and downstream homology arms (lane 2) of the Rv3671c gene. The results were analyzed by agarose gel electrophoresis. b): The digestion results of recombinant phAE159 with *SrfI* and *PflMI*. We selected 3 colonies of recombined HB101 from LB agar with hygromycin B. After obtaining their recombinant phAE159-*ΔMarp*, the phagemid was digested by *SrfI* and *PflMI* and analyzed by agarose gel electrophoresis (lane1-3).

### 3.2 Preparation of high-titer phage lysate

Recombinant phagemid was transfected into Mycobacterium smegmatis mc^2^155 competence by electroporation, and the plaques (shown in Figure 4a) were picked up to prepare high-titer phage lysate. The phage was amplified in M. smegmatis mc^2^155 and infected *M. bovis* for homologous recombination. The colonies of *M. bovis* were selected and detected by primers of LYZ and RYZ. As a final stage, the PCR results were closely examined by agarose gel electrophoresis as Figure 4b and showed that the hygromycin resistance gene successfully replaced the *Rv3671c* in ΔMarp mutant, and the results of western blotting (Figure 4c) proved that there was no Marp expression in ΔMarp mutant, indicating that ΔMarp strain was successfully constructed.

**Figure 4.**
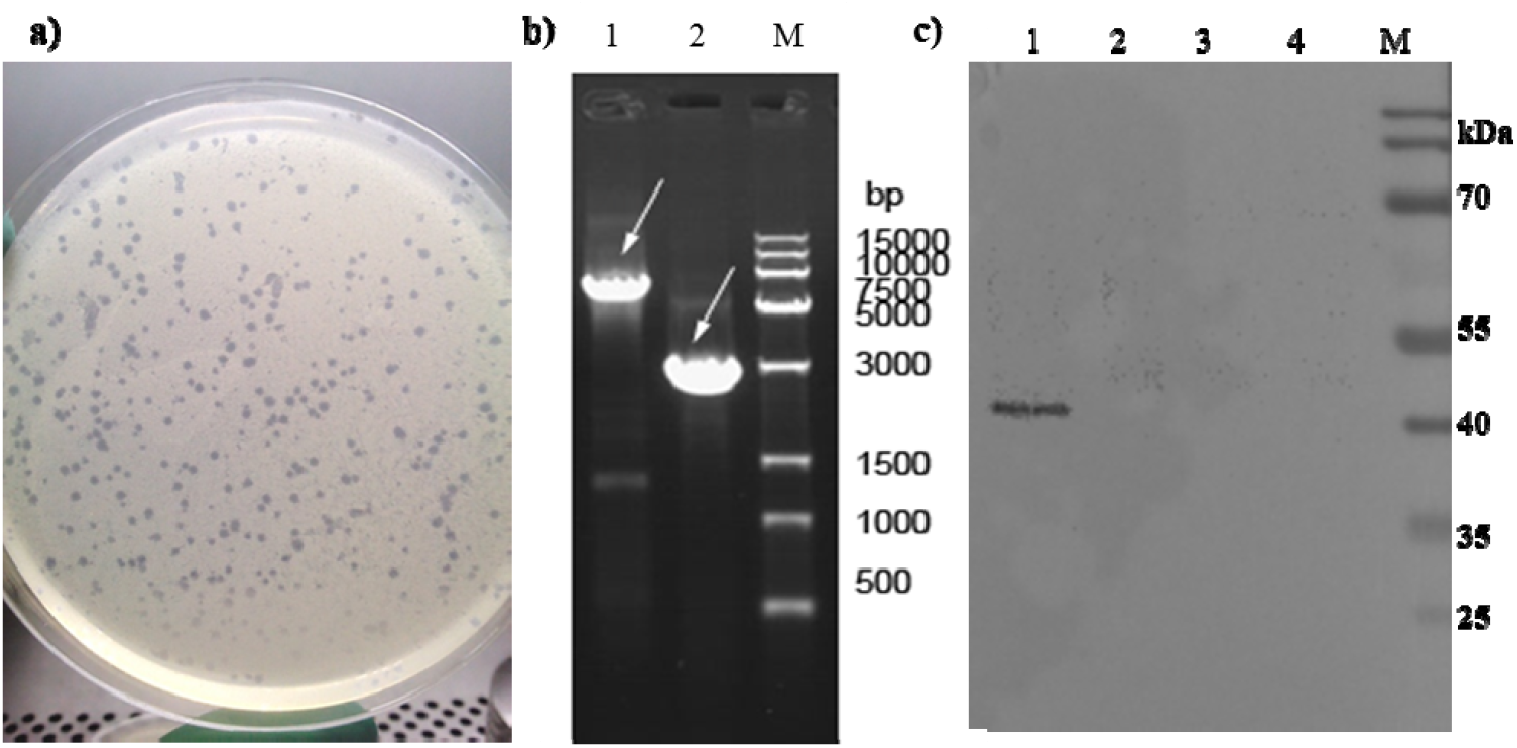
Identification of ΔMarp mutant. a): Plaque formed by the constructed bacteriophage. Recombinant phagemid was transfected into *M. smegmatis* mc^2^155 competence by electroporation, and the plaques will show up like above. b): Identification results of ΔMarp mutant by PCR. The colonies of *M. bovis* and wild-type AN5 were selected and detected by primers of LYZ and RYZ. The white arrow indicates the target DNA fragment. Lane 1 showed the DNA ladder that *Rv3671c* was replaced with *sacB* gene cassette in the mutant clone; lane 2 showed the original target *Rv3671c* gene DNA fragment in wild-type AN5. It means that the hygromycin resistance gene successfully replaced the *Rv3671c* in ΔMarp mutant. c): Identification of ΔMarp mutant by western blot. We selected 3 colonies of recombineering colonies from 7H10 agar with hygromycin B and extracted their total proteins. Using the Marp monoantibody, the Marp were detected in these colonies by western blot with the total proteins of wild-type An5 as a positive control. The results showed that Marp can be detected from wild-type AN5 (lane1) but there was no Marp expression in ΔMarp mutant (lane2-4).

### 3.3 Construction of Complemented Strain and overexpressing strain

PCR amplification helped to obtain the PCR products *Rv3671c*; PMV261 plasmids were digested with *EcoRI* and *Hind III* at the same time. After that, their digested products were ligated together, and then the conjunction was transformed into *E*.*coli* DH5α competent cells; the bacteria needed to be cultured on LB agar with kanamycin to select colonies whose plasmids were used to identify by PCR. The recommend plasmids were extracted and electroporated individually into the *ΔMarp* and An5; they were then cultured on 7H10 agar at 37°C for 4 weeks. The identification results about the selected colonies were obtained by PCR shown in Figure 5.

**Figure 5.**
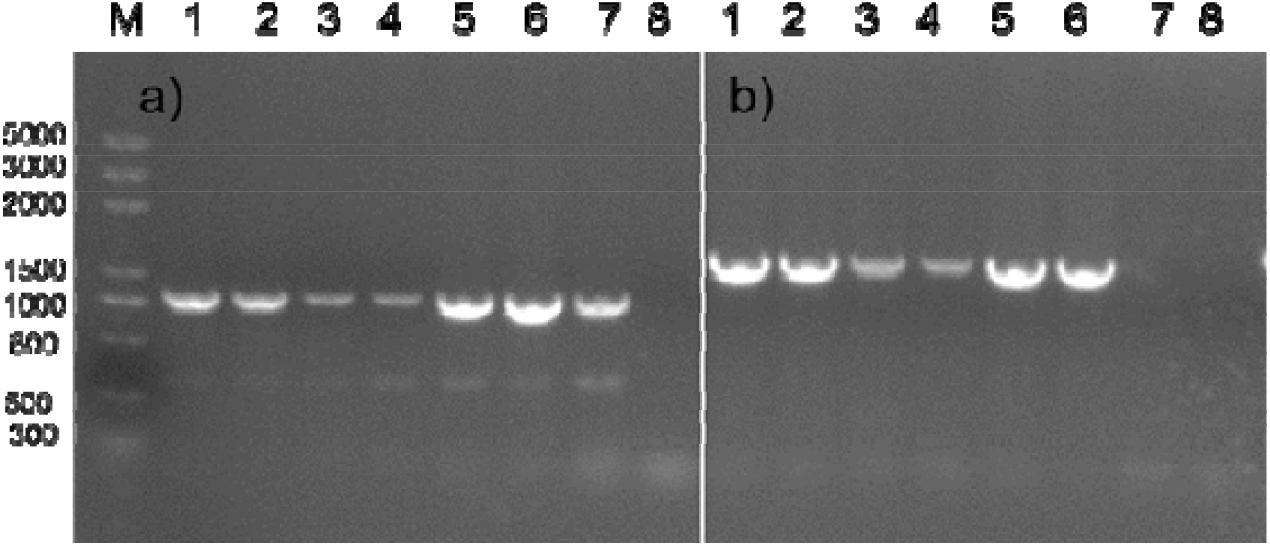
Identification results of ΔMarpComp and PmvRv3671 by PCR. The recombined bacteria (ΔMarpComp and PmvRv3671) were selected, and their DNA was extracted. Their DNA was detected by PCR with primers Marp (Figure 5a) or Pmv261 (Figure 5b); meanwhile, the An5 genome was used as the positive control, and ddH_2_O was used as the negative control. a) The PCR amplification results with Marp primers. It showed that the *Rv3671c* gene could be found in ΔMarpComp strains (lane 1-3), PmvRv3671 strains (lane 4-6) and An5(lane 7), and could not be found in the negative control (lane 8); it means that the PCR process was nonpolluted; b) The PCR amplification results with PMV261 primers, showing that Pmv261-Marp plasmid could be found in ΔMarpComp strains (lane 1-3) and PmvRv3671 strains (lane 4-6), and it could not be seen in An5 (lane 7) and ddH_2_O (lane 8).

### 3.4 Measurement of acid resistance and growth characters

All of the four types of bacteria were cultured in 7H9-pH6.6 and 7H9-pH5.0 mediums at 37°C (they all cannot grow at 7H9-pH 4.5), and their growth curves were measured during 14 days. The growth results of OD_600_ value and CFU about the four types of bacteria, in standard 7H9-pH6.6 medium, within 14 days are shown as Figures 6a and 6b. Figure 6a showed that the wild-type strain An5 presented the largest increase in OD_600_ values over time, and the other three strains were similar. Figure 6b represents the relationship between the number of living bacteria and the change of time. As it can be seen, the PmvRv3671 has more CFU than the other three. To sum up, the number of living bacteria of PmvRv3671 is 10^8.55^ on the 14^th^ day, 5 times more than that of *ΔMarp*. It suggested that the deletion of gene *Rv3671c* might weaken the growth ability of the strain in a neutral environment.

**Figure 6.**
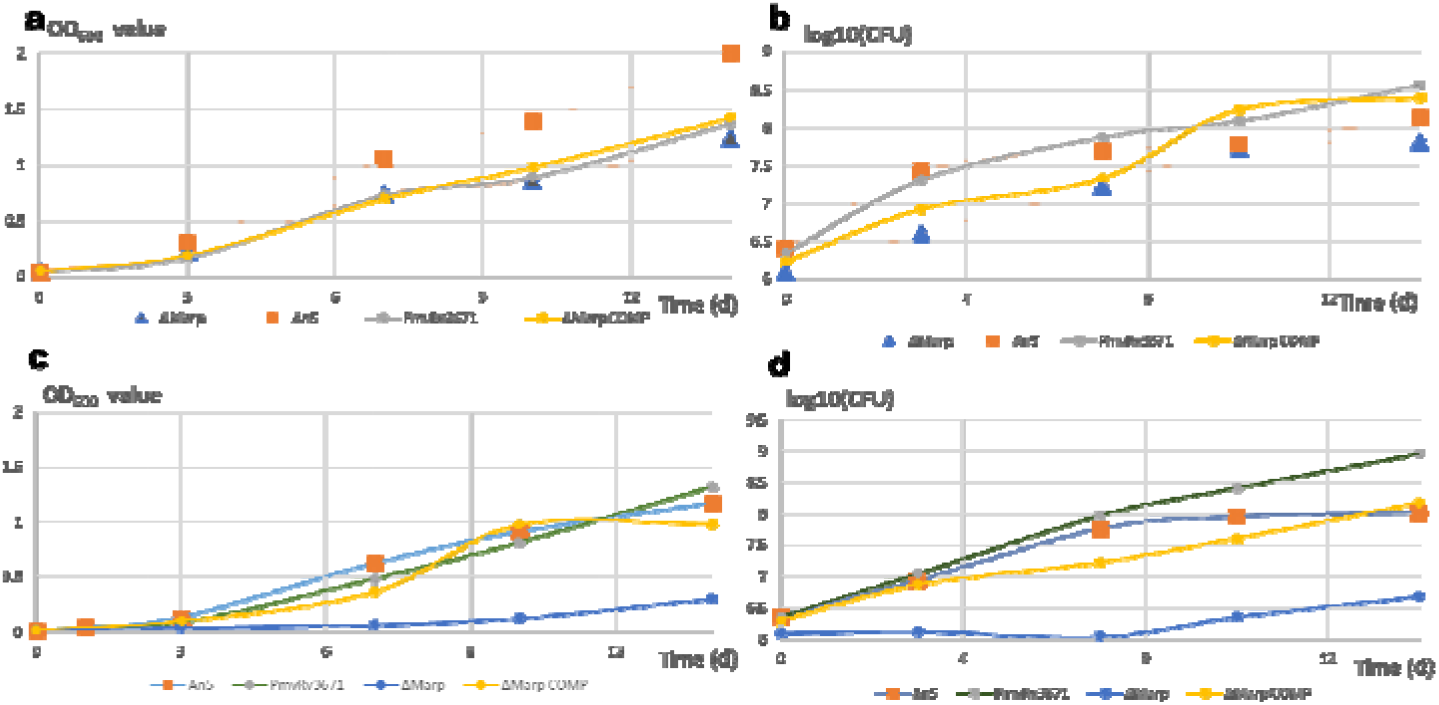
Growth results of An5, *ΔMarp*, ΔMarpComp, and PmvRv3671 respectively at pH 5.0 and pH 6.6 7H9. All of the four types of bacteria were cultured in 7H9-pH6.6 and 7H9-pH5.0 medium at 37°C, and their growth curves were measured during 14 days. (a) OD_600_ value of An5, *ΔMarp*, ΔMarpComp, and PmvRv3671 at pH 6.6 in 7H9; (b) Quantification of CFU of An5, *ΔMarp*, ΔMarpComp, and PmvRv3671 at pH 6.6 in 7H9; (c) OD_600_ value of An5, *ΔMarp*, ΔMarpComp, and PmvRv3671 at pH 5.0 in 7H9; (d) Quantification of CFU of An5, *ΔMarp*, ΔMarpComp, and PmvRv3671 at pH 5.0 in 7H9. Bacterial input was 0.5× 10^7^ CFU ml^−1^.

The growth curve of OD_600_ value and CFU of the four types of bacteria, in 7H9 at pH 5.0 medium, within 14 days are exhibited in Figures 6c and 14d. Figure 6c shows that the OD_600_ value of *ΔMarp* on the 14^th^ day was 0.30, the lowest value among these four strains. The OD_600_ value of An5, ΔMarpComp, and PmvRv3671 were 1.17, 0.98, and 1.32, respectively. From the comparison of the count of living bacteria in Figure 6d, it can be seen that the PmvRv3671 had the highest value of log_10_ CFU/mL – 8.95 in 7H9-pH5.0, almost 0.4 logs more than that in 7H9-pH6.6. The log_10_ CFU/mL value of ΔmarpComp and An5 are similar – 8.16 and 8.01 – either of them is more 2.0 log than the value 6.68 of *ΔMarp*. This indicates that the *Rv3671c* gene affected the growth of *Mycobacterium bovis* in 7H9-pH 5.0 medium.

In summary, PmvRv3671 had more living bacteria in an acidic environment than a neutral environment. Moreover, the *ΔMarp* cannot maintain the ability to grow in an acidic environment like wild-type strain, so it suggested that, due to the existence of gene *Rv3671c, M. bovis* could grow in an acidic environment like in the neural environment. In other words, *Rv3671c* may participate in the acid-resistant process of *M. bovis* and the process of maintaining the intracellular neutral environment under acidic environment.

### 3.5 Determining the different acid resistance of ΔMarp, ΔMarpComp, PmvRv3671, and An5

For this step, the four strains were cultured in a biochemical incubator at 37°C at pH 4.5 and pH 6.6 in pcit-ty maintenance buffer. We have measured their survival rates for 14 days. As can be seen from Table 2, on the 5^th^ day, the survival rates of *ΔMarp*, ΔMarpComp, An5, and PmvRv3671 in the acidic (pH4.5) environment were 12.94%, 25.65%, 21.02%, and 109.4%, respectively; meanwhile, the survival rate of PmvRv3671 in the acidic environment was even better than that in a neutral environment.

**Table 2.** Survival rates of *M. bovis* An5, *ΔMarp*, ΔMarpComp, and PmvRv3671 respectively at pH 4.5 and pH 6.6 maintenance solutions (pcit buffer).

|  | 5d |  |  | 14d |  |  |
| --- | --- | --- | --- | --- | --- | --- |
|  | pcit-6.6 | pcit-4.5 | pcit-4.5/pcit-6.6<br># | pcit-6.6 | pcit-4.5 | pcit-4.5/pcit-6.6<sup>#</sup> |
| An5 | 73.03%±2.15<br>%<sup>a</sup> | 21.02%±1.10<br>%<sup>a</sup> | 28.8%±2.61%<sup>aA</sup> | 6.03%±0.23%<sup>a</sup> | 0.68%±0.02<br>%<sup>a</sup> | 11.26%±0.5%<sup>a</sup><br>B |
| ΔMarp | 69.97%±6.06%<sup>a</sup> | 12.94%±0.29%<sup>c</sup> | 18.56%±0.7%<sup>b</sup><br>A | 69.69%±3.03<br>%<sup>c</sup> | 0.09%±0.00<br>%<sup>b</sup> | 0.13%±0.005<br>%<sup>bB</sup> |
| ΔMarp<br>Comp | 83.33 %±4.41%<sup>b</sup> | 25.65%±1.57%<sup>a</sup> | 30.8%±3.25%<sup>aA</sup> | 12.673%±0.44<br>%<sup>b</sup> | 0.72%±0.10<br>%<sup>a</sup> | 5.66%±1.36%<sup>a</sup><br>B |
| PmvRv36<br>71 | 67.783%±2.94%<sup>a</sup> | 109.403%±4.45<br>%<sup>b</sup> | 161.4%±11.37<br>%<sup>cA</sup> | 5.113%±0.48<br>%<sup>a</sup> | 1.70%±0.27<br>%<sup>c</sup> | 33.29%±9.2%<sup>c</sup><br>B |

Note: The superscript ‘#’ represents the survival rate under pcit-4.5 divided by survival rates under pcit-6.6, used to correct the effect of pcit buffer, and represents the survival rate under acidic environment. The different superscript ‘a~c’ means the difference in the survival rates within this column is highly significant ( $P<0.01$ ); meanwhile, the different superscript ‘A~B’ means the difference in the survival rates of the same bacteria between 5<sup>th</sup> and 14<sup>th</sup> day is highly significant ( $P<0.01$ ). Data are means ± s.d. of triplicate cultures and represent two or more independent experiments.

Only after dividing the survival rate of each strain at pH 4.5 by their own survival rate of pH 6.6 in pcit buffer, the corrected survival rate could be calculated. It can be used to evaluate the resistance to the pressure from the acid environment without the effect of pcit buffer. The adjusted values revealed that the survival rate of the PmvRv3671 (161.4%) in pH4.5 environment was very significantly higher than that of the ΔMarpComp (30.8%) and An5 (28.8%), both significantly higher than *ΔMarp* (18.56%).

According to statistic analysis results showed in Table 2, on the 14^th^ day, the survival rate of *ΔMarp* in the pcit-6.6 is 69.97%, higher than that of the other three strains. Next is the ΔMarpComp (12.67%), more inflated than the An5 (6.03%) and PmvRv3671 (5.11%) in the neutral environment. Contrarily, the survival rate of PmvRv3671 in the pcit-4.5 is 1.7%, also higher than that of ΔMarpComp (0.72%), An5 (0.68%) and *ΔMarp* (0.09%). The survival rates of *ΔMarp* (0.09%) is the lowest one among these four strains on 14^th^ day. Without the influence of the maintenance solution, the survival rate of PmvRv3671 (33.29%) is still more advantageous than An5 (11.26%), ΔMarpComp (5.66%) and *ΔMarp* (0.13%).

In summary, we found that the survival rate of the PmvRv3671 is the lowest one in pcit-6.6 and the highest one in pcit-4.5 among the four strains. On the other hand, *ΔMarp* is particularly stable in the environment of pcit-6.6, even on the 14^th^ day, the survival rate was the same as the 5^th^ day (69.97%). Besides, comparing the corrected data of the four strains, the acid resistance capacity of each four strains on the 14^th^ day has decreased significantly than that on the 5^th^ day. These indicated that the presence or absence of the *Rv3671c* gene made a notable influence on the survival rate of *M. bovis* in a neutral and acidic environment, and *ΔMarp* remained part of acid resistance, but the ability of *ΔMarp* also became weaker with the time extended.

### 3.6 The different growth rates of ΔMarp, ΔMarpComp, PmvRv3671, and An5 in Raw264.7

After 48 hours of infection, most bacteria have been colonized into Raw264.7. The number of bacteria in cells was measured at 48 hours, 96 hours, and 144 hours after infection to calculate the bacterial growth rate. From Figure 7a, few distinct growth characteristics of these four strains could be spotted. From 48h to 96h after infection, the growth rate in the resting Raw264.7 was significantly faster than in cells after IFN-γ stimulation; *ΔMarp* almost didn’t grow in the unstimulated Raw264.7 at 96h-144h. The growth rate of the other three strains between 96h to 144 hours was highly significantly faster than in the other periods. Also, between 96 and 144 hours, the four strains were growing significantly faster in the resting Raw264.7 than in stimulated cells.

**Figure 7.**
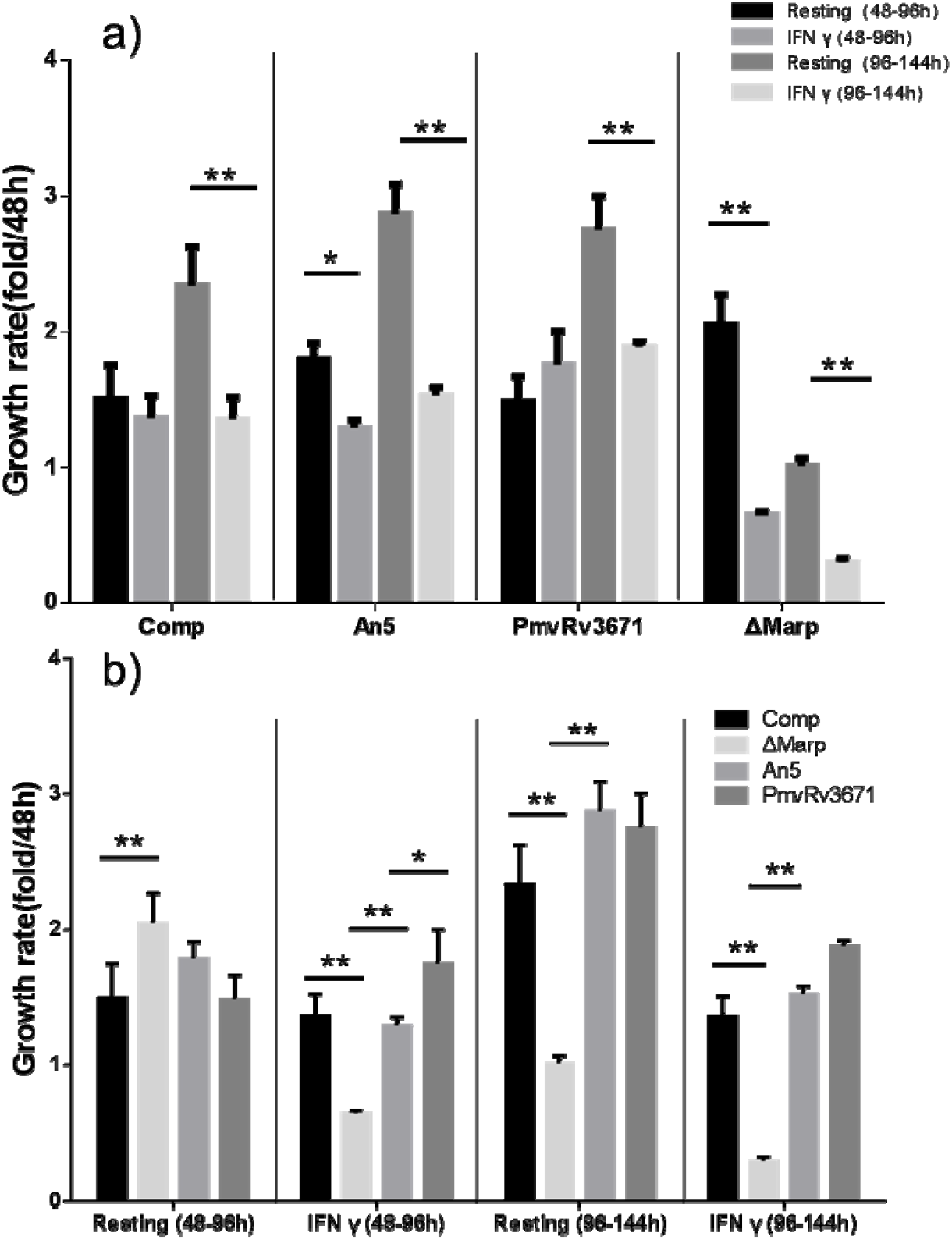
Growth rates of *M. bovis* An5, *ΔMarp*, ΔMarpComp, and PmvRv3671 respectively in resting Raw264.7 and Raw264.7 stimulated by IFNγ. a) Growth rates of these four strains in resting Raw264.7 and Raw264.7 stimulated by IFNγ in different periods; b)Comparison of the growth rates of these four strains in resting Raw264.7 and Raw264.7 stimulated by IFNγ. ‘<u>*</u>’or’<u>**</u>’ means the differences of the growth rates between the two groups is significant (*p*<0.05) or highly significant (*p*<0.01). Data are means ± s.d.of triplicate cultures and represent two or three independent experiments. In some panels, error bars are too small to be seen.

Some prominent growth characteristics could be seen in Figure 7b. The growth rate of *ΔMarp* was significantly higher than that of the ΔMarpComp at 48h-96h after infection in unstimulated Raw264.7. In Raw264.7 after IFNγ stimulation, the growth rate of the *ΔMarp* is very significantly slower than that of the other three. The growth rate of PmvRv3671 and An5 are similar in resting Raw264.7 or with IFNγ stimulation.

In summary, we found that as time goes on, the growing rates of ΔMarpComp, PmvRv3671, and An5 all are higher between 96 and 144 h than between 48 and 96 h in resting Raw264.7 cells or Raw264.7 cells with IFNγ stimulation, but the situation of *ΔMarp* is contrasting. This indicated that the presence or absence of the *Rv3671c* gene made a notable influence on the growth rate of *M. bovis* in Raw264.7 cells.

## 4 Discussion

Mycobacterium tuberculosis is a typical intracellular pathogen, which can survive in extreme conditions such as hypoxia, nitric oxide killing, and nutritional deficiencies in macrophages, and suppress the host immune response through multiple escape mechanisms to avoid clearance by the body (Mena Cimino et al., 2012; Naffin-Olivos et al., 2014; Philips and Ernst, 2012; Walburger et al., 2004). Although some are killed under those changing environmental conditions, the escaped MTB could still mainly parasitize in macrophages and replicate or replicate into a stage of dormancy rendering itself extremely resistant to host defence (Mena Cimino et al., 2012). In a word, the pathogenic mechanisms in hosts are very complicated and the *Rv3671c* gene is essential for the process of acid resistance. *M. bovis* is very similar to MTB but has a wider host, and it has been confirmed that this gene also presented in *M. bovis* with our previous experiments, but whether there are other biological functions like growth involved remains to be a mystery to be explored. To further verify the functions and biological characteristics of the gene *Rv3671c* in *M. bovis*, the *Rv3671c* knockout strain was prepared using the method as previously (Jain et al., 2014), and the shuttle vector was utilized to construct the complemented strain and overexpression strains.

According to the measurement results of the growth curves of these four strains in pH6.6 7H9, the *ΔMarp* could grow normally, which is similar to the results reported previously (Botella et al., 2017), but the CFU of PmvRv3671 on 14^th^ day is more than five times with the CFU of the *ΔMarp*. This suggested that the deletion of *Rv3671c* in *M. bovis* could affect its growth, but it is not an essential gene for *M. bovis* growing in standard 7H9 medium. However, when the experiment overexpressed *Rv3671c* gene in *M. bovis*, PmvRv3671 showed a stronger growth performance in a nutritious medium, indicating that the gene might enhance the related metabolisms of *M. bovis*. Although the OD_600_ value of PmvRv3671 on the 14^th^ day was lower than that of the An5 strain, it might be related to the change in color of the bacteria, and this characteristic could be observed in PmvRv3671.

From the growth curve measurement results of the four strains cultured in pH6.6 7H9, it was found that the growth results of PmvRv3671 also outperformed An5 and this point was amplified in acidic nutrient medium (pH4.5-7H9). Meanwhile, *ΔMarp* could not maintain the growing ability in that acidic medium. The above two points both indicated that the *Rv3671c* gene infected the growth of *M. bovis*, especially in an acidic nutrient medium. Vandal (Vandal et al., 2008) previously performed another experiment on the Mtb H37Rv at 7H9-Tw-4.5 medium, which turned out to be that the inoculated Mtb was almost completely killed on the 15th day. After that, they proposed Tween-80 might release free fatty acids to kill Mtb, so Tween-80 was replaced by nonhydrolyzable tyloxapol as the dispersing agent, but the change still failed to observe the growth of Mtb in pH4.5 7H9. This exhibited that the wild-type *M. bovis* An5 share similar acid-resistance mechanisms like Mtb, because the *M. bovis* An5 cannot grow in pH4.5 7H9 either. In addition, PmvRv3671 grew better in acidic environment (pH5.0 7H9) than the neutral environment, which suggested that *Rv3671c* gene plays a more important role in acidic environments; it might be due to the acidic environment trigger more pathways by activating *Rv3671c* gene and its related downstream, like ripA response for cell elongation and chain formation (Botella et al., 2017); Flannagan also believes that acidic conditions will activate the degradation of bacterial lipids and proteins (Flannagan et al., 2009). For instance, *Escherichia coli* (*E. coli*) could use amino acid decarboxylase/antiporter systems that utilize glutamate, arginine, and lysine as substrates, to increase its cytoplasmic pH (Foster, 2004), and Marp, a serine protease, might work in a similar way to help the growth of mycobacteria.

By observing the survival rate results of the four bacteria in pH 4.5 and pH 6.6 maintenance solutions (pcit buffer), it can be seen that the rate of *M. bovis* in neutral and acidic environments both decrease as time goes on, except *ΔMarp* in neutral environment. PmvRv3671 presented the lowest survival rates on both the 5^th^ and the 14^th^ day when culturing in the pcit pH 6.6 environment, but no significant difference with An5 was observed. Notably, *ΔMarp* was extremely stable in the pcit pH 6.6 environment, and even on the 14th day, the survival rate was the same as the 5^th^ day (69.97%). This indicates that the *Rv3671c* gene does not promote the survival of *M. bovis* in the neutral maintenance environment; contrarily, *ΔMarp* can survive longer in a neutral environment and it might since the deletion of the *Rv3671c* gene which could slow parts of the metabolisms. Previously it has been proved that Marp cleaves the peptidoglycan hydrolase RipA to activate RipA in acid environment (Botella et al., 2017). When Marp was knockout in *ΔMarp*, the failure of RipA processing leads to cell extension and chain formation, which is a sign that the separation of progeny bacteria stops. The lack of interaction with RipA in *ΔMarp* resulted in the activity of RipA is greatly reduced. This inhibited the bacterial growth rate, which is why it could be observed that PmvRv3671 grows faster than An5 and An5 also grow faster than *ΔMarp*. This also explains why the deleted strain in the neutral maintenance solution could be so stable even on the 14^th^ day. Surprisingly, the proteomic comparison exhibited that the ATP γ chain in the deleted strain was significantly more than that in the wild strain (Small, 2013). It further suggests that the *ΔMarp* use and consume ATP less efficiently and the deletion of Marp in *M. bovis* might slow down its energy metabolisms.

On the other hand, the survival rate of PmvRv3671 in pH 4.5 maintenance buffers is even higher than that in pH 6.6 maintenance buffer, providing a similar outcome to the results culturing it in 7H9 medium. Comparing the corrected survival rates, it can be found that after removing the influence of the maintenance solution, the acid resistance of them on the 14^th^ day all were significantly decreased than that on the 5^th^ day. Apparently, the *ΔMarp* still remain part of acid resistance; even without Marp there are still other acid-resistance mechanisms like the predicted magnesium transporter MgtC (Buchmeier et al., 2000); outer membrane protein OmpATB (Song et al., 2011); a two-component regulator PhoPR (Rohde et al., 2007); a gene locus regulated by PhoPR in response to acid aprABC (Abramovitch et al., 2011) in mycobacteria. Whatever, the comprehensive acid resistance of *M. bovis* still becomes weaker in acid maintenance solution as time goes on.

According to the growth rate of these four strains in Raw264.7 macrophages, it can be stated that mature macrophages presented a stronger killing capacity on mycobacteria. The growth rates of *M. bovis* An5,ΔMarpComp, and PmvRv3671 all are greater than 1.0 in mature macrophages, which indicates that these three strains can grow or even further infected macrophages; *ΔMarp*, however, is dying and cannot maintain the infection state for a long time in mature macrophages, even in the resting macrophage in the 96-144h. Therefore, it can be seen that the deletion of the *Rv3671c* gene prevents *M. bovis* from forming a long-term infection in mature macrophages, indicating that the *Rv3671c* gene is essential for the survival of *M. bovis* in mature macrophages like previous in MTB (Vandal et al., 2008). In addition, PmvRv3671 exhibited a faster growth rate than wild-type strains in mature macrophages like it did in 7H9 at pH 5.0. Besides, comparing their growth rates in macrophage with IFN-γ stimulated, it proved that mature macrophage can not eliminate all intracellular bacteria. However, comparing their growth in unstimulated Raw264.7 cells, even if mature macrophages can not eliminate all intracellular bacteria, they remain the inhibitory ability to intracellular mycobacteria.

Tuberculosis is an important infectious disease with large number of deaths caused by a single pathogen in the world and is also considered the leading cause of death after AIDS (WHO, 2019).

The main pathogenic bacteria of tuberculosis are Mycobacterium tuberculosis (Mtb) and Mycobacterium bovis (*M. bovis*). Mechnikov indicated that macrophages would kill most of the ingested microorganisms by acidifying its interior, but Mtb could use its waxy cell wall to resist this acidification (Mechnikov, 1988). According to previous reports, when Bacille Calmette-Guérin (BCG), an attenuated variant of *M. bovis*, was exposed to pH 5.5 in vitro, it ceased replicating and Mtb continued to divide slowly unless the pH was lowered to 4.5. With the addition of 0.5 mM nitrite to 7H9 medium at pH 5.5, Mtb would also stop replicating. However, it is in a balanced state, where the Mtb could survive for several days, and the bacterial can return to a normal growth rate after replacing the medium with standard 7H9 (Bryk et al., 2008; Gold et al., 2012; Vandal et al., 2008).

Based on that, Vandal used the acidic environment to screen 10,100 MTB transposon mutants and found a mutant that cannot maintain its intracellular pH homeostasis under a pH 4.5 environment (Vandal et al., 2008). Because the gene *Rv3671c* product played an essential role in the acid resistance of Mtb, it is named as Mycobacterial acid resistance protease (Marp). Marp was identified as a transmembrane serine peptidase with a protease domain located in the periplasm, as confirmed by a prudent analysis of its mutations, homology modeling, crystallography, and substrate profiling (Biswas et al., 2010; Small et al., 2013; Vandal et al., 2008).

In summary, culturing the four bacteria grew in 7H9 medium at pH 5.0 or pH 6.6 and maintenance buffer at pH 4.5, the lived number of over-expressing strain PmvRv3671 is significantly more than wild-type strain *M*.*bovis*, ΔMarpComp and *ΔMarp* on the 14^th^ day. After removing the effects of citrate solution, it can be found that the acid resistance abilities of them all are significantly lower on the 14^th^ day than that on the 5^th^ day. Using them infected Raw264.7 macrophages with IFNγ stimulation, the growth rate of the PmvRv3671 is faster than An5, *ΔMarp* and ΔMarpComp. In conclusion, the gene *Rv3671c* in *M. bovis* is not only related to acid resistance but also affected the growth of *M. bovis*, even the metabolisms of energy.

